# The reef-building coral *Galaxea fascicularis*: a new model system for coral symbiosis research

**DOI:** 10.1101/2022.06.02.494472

**Authors:** Giulia Puntin, Jamie Craggs, Róisín Hayden, Kara E. Engelhardt, Shelby McIlroy, Michael Sweet, David M. Baker, Maren Ziegler

## Abstract

Reef-building corals owe their evolutionary success to their symbiosis with unicellular algae (Symbiodiniaceae). However, increasingly frequent heat waves lead to coral mass-bleaching events and pose a serious threat to the survival of reef ecosystems. Despite significant efforts, a mechanistic understanding of coral-algal symbiosis functioning, what leads to its breakdown and what can prevent it, remains incomplete. The main obstacles are low amenability of corals to experimental handling and, owing to its obligatory nature, the difficulties of manipulating the coral-algal association. Indeed, many studies on the symbiotic partnership are conducted on other cnidarian model organisms and their results may therefore not be fully transferable to tropical reef-building corals. Here, we identify the tropical stony coral species *Galaxea fascicularis* as a novel candidate coral model system. Individual polyps of this species can be separated, enabling highly replicated genotype studies, and are well suited to experimental investigation of the symbiosis as they can be easily and effectively rid of their algal symbionts (bleached). We show that bleached adult individuals can reestablish symbiosis with non-native symbionts, and we report the completion of the gametogenic cycle *ex-situ*, with the successful spawning in aquaria over multiple years. These achievements help overcome several of the major limitations to direct research on corals and highlight the potential of *G. fascicularis* as an important new model system for investigations of symbiosis functioning and manipulation.

## Introduction

Reef-building corals form obligatory endosymbiotic association with unicellular algae of the family Symbiodiniaceae. This association is key to their evolutionary success, but it is also at the heart of corals’ susceptibility to global climate change, which manifests in coral bleaching - the breakdown of the coral-algal symbiosis (Hoegh-Guldberg 1999). Bleaching is mostly driven by marine heatwaves which are predicted to worsen, causing the loss of virtually all coral reefs by the end of this century (van Hooidonk et al. 2016).

The interaction between heat stress and bleaching has been studied for more than 30 years (Gates et al. 1992; McLachlan et al. 2020), but a detailed understanding of the mechanisms underlying coral bleaching is still missing. Progress in coral symbiosis research is hampered by two main aspects. First, corals are challenging to maintain in aquarium settings as they tolerate only a narrow range of environmental conditions, i.e., bleaching can be triggered by just 1-2 °C increases above summer mean temperatures (Glynn and D’Croz 1990). Second, the obligatory nature of the coral-algal association makes it particularly challenging to physically and functionally separate the partners, an approach often necessary to unravel the mechanisms underlying symbiosis breakdown and what could prevent it (Weis et al. 2008; Voolstra 2013).

Model organisms are integral for understanding fundamental biological principles and symbiotic cnidarians have successfully been used as “coral models”, advancing our understanding of coral bleaching and holobiont functioning (Weis et al. 2008). These model organisms share important traits with corals, such as being cnidarians that associate with microalgae, yet they also lack other features that normally represent obstacles to research work such as the calcium carbonate skeleton. In addition, the ability to study the animal host and its algal symbiont in isolation is more easily achieved with facultatively symbiotic organisms such as *Hydra* spp., Aiptasia (*Exaiptasia diaphana*), and *Astrangia poculata* (Dimond and Carrington 2007; Weis et al. 2008; Galliot 2012). Cnidarian model organisms allow the study of basal or shared traits or phenomena at a speed that would not be possible with reef-building corals, but testing on the latter remains necessary for the understanding of coral-specific or ecologically relevant aspects. The obligatory symbiosis with Symbiodiniaceae in reef-building corals has important biological implications (Falkowski et al. 1984; Hoegh-Guldberg 1999), similar to the calcification process that is deeply intertwined with coral physiology and ecology (Gattuso et al. 1999). It is therefore important to establish a “true” tropical reef-building coral model species.

The richness of the scleractinian taxon offers a vast array of species to choose from in the quest for a suitable candidate coral model species. These can be evaluated for their tractability (as amenability to experimental work) considering several aspects which, as proposed by Puntin et al. (2022) should include: 1) pre-existing knowledge: baseline information of the organism’s biology is necessary to interpret and contextualize results; 2) compatibility with aquarium rearing: the possibility to maintain the organism and preferably to complete its life cycle in the lab is essential to increase its availability, reduce confounding effects such as unknown life history, improve reproducibility, and lower the pressure on threatened wild populations; 3) amenability to symbiosis manipulations is necessary to help unravel complex coral-algal functional interactions, and can be broken down into the organism’s suitability to be rendered aposymbiotic (bleached), to be maintained aposymbiotic for a certain time, and to then be re-infected with Symbiodiniaceae.

We have identified the coral species *Galaxea fascicularis* (Linnaeus, 1767) (clade: Complexa, family: Euphyllidae) as a promising candidate coral model system. The species is relatively common across the Indo-Pacific region (Veron et al. 2016), where the phylogeny of the genus *Galaxea* is geographically well resolved (Wepfer et al. 2020b). It is also one of the more commonly used species in heat-stress experiments (McLachlan et al. 2020) and in studies on coral calcification (e.g., Marshall and Clode 2004; Al-Horani 2005). Importantly, other community resources such as an annotated draft genome are available (Liew et al. 2016; Niu et al. 2016, http://www.gfas.reefgenomics.org/). In addition, *G. fascicularis* is known for its tolerance to stressors as it is “largely unaffected by bleaching” in the field (Marshall and Baird 2000) and for being easy to grow and propagate in aquaria (Pavia and Estacion 2019). Further, its morphology provides additional practical advantages with relatively big polyps, which can be easily isolated (Pavia and Estacion 2019). This conveniently allows reduction of the complexity of the organism from colonial to individual, enables high clonal replication, and facilitates visualization (Al-Horani et al. 2003; Marshall and Clode 2004). Its association with Symbiodiniaceae is relatively well characterized (Huang et al. 2011; Wepfer et al. 2020a). Furthermore, the species’ reproductive mode and spawning patterns in the wild are well documented (Babcock et al. 1986; Harrison 1988; Keshavmurthy et al. 2012). However, its amenability to symbiosis manipulation, to experimental handling of bleached individuals, and the ability to complete gametogenic cycles and spawning in closed aquaria have not yet been described.

The overarching aim of this study was to explore the potential of *Galaxea fascicularis* as a coral laboratory model. Specifically, we investigated: 1) the feasibility to render polyps aposymbiotic by eliminating Symbiodiniaceae (bleaching), 2) the amenability to experimental manipulation of symbiotic and aposymbiotic polyps in a simplified system, 3) the re-establishment of the symbiosis with different species of Symbiodiniaceae, and 4) the possibility to induce full gametogenic cycles *ex situ*, with subsequent spawning under aquarium conditions. These aspects are central to overcoming the two main obstacles to coral research work: low tractability of corals and their complex nature.

## Materials and methods

### Coral collection and long-term aquarium rearing

Colonies of *Galaxea fascicularis* were collected from three geographical locations: the Red Sea, Hong Kong, and the Great Barrier Reef (Fig. 1). To increase comparability, rearing conditions were adjusted to be similar between labs, with few exceptions pertaining to the particular experimental needs or feeding regime, which was based on the long-term rearing arrangements in each facility (Tab. S1). Red Sea colonies were collected at 9-13 m depth at the North-Eastern protected end of the reef “Al Fahal” in the central Saudi Arabian Red Sea (22.3054°, 38.9655°) in March 2019, and transported to the Ocean2100 aquarium facility at Justus Liebig University Giessen (Germany) where they are part of the live collection (CITES permit 19-SA-000096-PD). In the aquarium system, light was provided by white and blue fluorescent lamps with a light:dark cycle of 12:12 h at 130 −160 μmol photons m^−2^ s^−1^, which approximates light condition at the collection site (Ziegler et al. 2015). Salinity was maintained around 35 and temperature at 26 °C. Colonies were fed daily with a combination of frozen copepods, *Artemia*, krill, and *Mysis*.

**Fig. 1.**
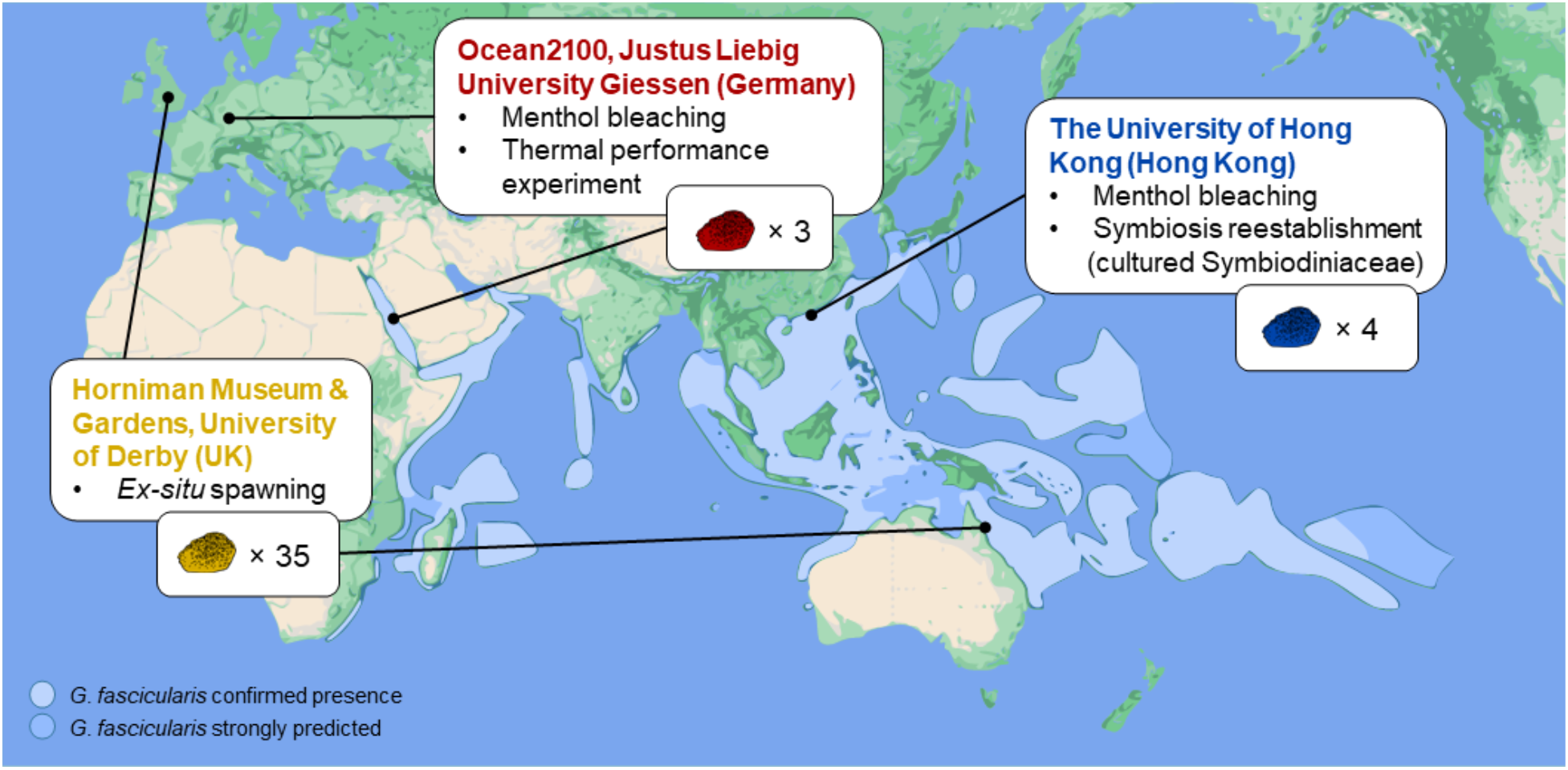
Summary of *G. fascicularis* coloniy origins, participating labs, and experiments described in this study. Colonies were collected from three distant locations (Red Sea, Hong Kong, and the Great Barrier Reef) and separately employed in different experiments (menthol bleaching, thermal performance, symbiosis reestablishment, and *ex situ* spawning). Map modified from Corals of the World (www.coralsoftheworld.org, Veron et al. (2016)).

Hong Kong colonies were collected from 5 m (max. depth) from Crescent Island (22.5308°, 114.3150°) in November 2020 and transported to the University of Hong Kong, where they were fragmented and acclimated to aquarium conditions for one month. The aquarium system consisted of 6-L acrylic tanks placed within a Plant Growth Chamber (Panasonic MLR-352H-PA). Each tank held eight ~3 cm-fragments and was fitted with a submersible pump (Atman AT301) and a small water filter (Shiruba PF120). The fragments used in symbiosis reestablishment were fed twice-weekly with a powdered blend of marine plankton (ReefRoids, Polyp Lab) followed by a partial water change. Light, salinity, and temperature conditions were consistent with those maintained in the Ocean2100 facility.

For *ex situ* spawning, 35 colonies (10.1 cm ± 2.5 (mean ± sd)) were collected from Arlington Reef, Great Barrier Reef, Australia (−16.7000°, 146.0500°) in September 2019, and transported to the Horniman Museum and Gardens (United Kingdom; CITES permit 585319/01) with the ‘inverted submersion method’ as described by Craggs et al. (2018). Colonies were distributed into four coral spawning systems (www.coralspawninglab.org) that replicated the natural environmental parameters (seasonal temperature, photoperiod, solar irradiance, and lunar cycle) required to stimulate reproduction (Tab. S1). Aquarium design and coral husbandry protocol was based on the mesocosms described by Craggs et al. (2017), and the seasonal temperature profile were based on a non-sequential eight year average (1998 - 2017) from Moore Reef (−16.8667°, 146.2334°’) (http://data.aims.gov.au/aimsrtds/datatool.xhtml?site=931&param=water%20temperature) (Fig. S7). Colonies were fed daily on a mix of phytoplankton (*Tisochrysis lutea*, *Chaetoceros calitrans* and *Rhodomonas salina*) and zooplankton (newly hatched *Artemia salina* nauplii, frozen red plankton, rotifers, and lobster / fish eggs) (Tab. S1). During feeding the broodstock tank was isolated from the filtration for two hours to aid prey capture and uptake.

### Thermal performance experiment on single polyps

#### Preparation of single polyps

For the menthol bleaching and thermal performance experiment, we used single polyps from three Red Sea colonies each showing distinct coloration (Fig. S1). Replicate clonal polyps were isolated from each colony using an electric rotary cutter, mounted on coral glue (Grotech, CoraFix SuperFast), and allowed to heal for two weeks in the Ocean2100 aquarium facility (Schubert and Wilke 2018).

#### Menthol bleaching to remove algal symbionts

An equal number of polyps from each colony were randomly assigned to the control group (maintained in a symbiotic state), and the bleached group (chemically bleached with menthol). Menthol bleaching followed a protocol modified from Wang et al. (2012) and consisted of three days of treatment in 0.38 mM menthol solution in seawater, followed by a day of rest and a fourth day of menthol treatment. Each day the polyps were incubated in menthol for 8 h under light and stirring (Fig. S2). After, the polys were rinsed and kept in clean containers with air bubbling and stirring. During menthol treatment, the treated polyps were only exposed to filtered (1.2 μm) artificial seawater (FASW) to prevent exposure to Symbiodiniaceae. Following, all polyps (including, for consistency, the symbiotic group) were maintained in FASW.

To confirm bleaching, half of the polyps from each group (bleached and symbiotic) and each colony were visually assessed with a fluorescence stereomicroscope (Leica MZ16 F) 10 days after the termination of the menthol treatment. Representative pictures were taken under natural light (brightfield, light source: 2950 K) and under UV light with GF2 filters allowing visualization of the green fluorescent proteins and chlorophyll (Fig. 2a). Following the same bleaching protocol, menthol-bleached polyps were also documented using a compound epifluorescence microscope to reach higher resolution (Leica DM 5500B, TX2 filter) (Fig. 2b).

**Fig. 2.**
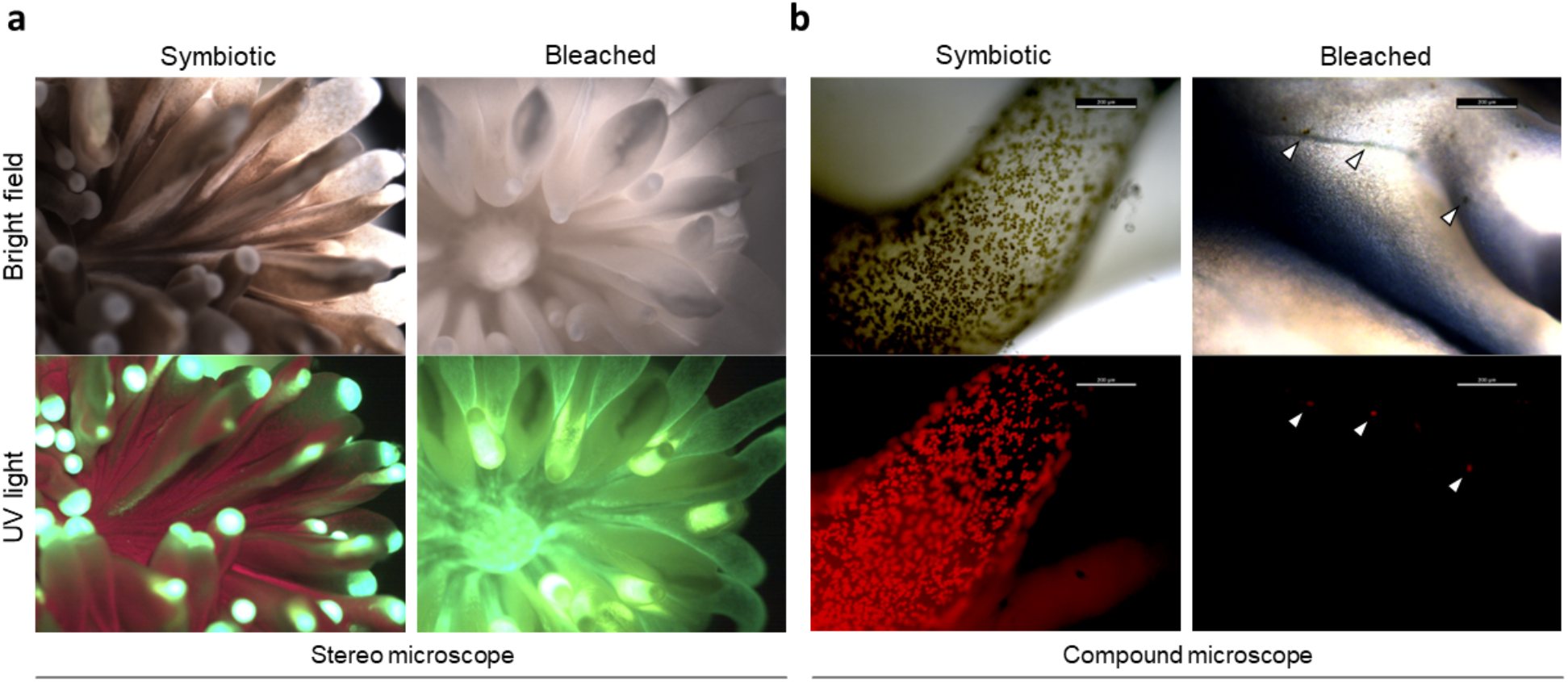
Microscopic comparison between symbiotic and menthol-bleached polyps. Representative stereo and compound micrographs in bright field and UV light, with filters for chlorophyll (red) and coral tissue (green) autofluorescence. **a** Symbiodiniaceae cells are abundant in symbiotic polyps while not detectable in bleached polyps. **b** At higher magnifications, few algal cells are still detectable in bleached polyps only at the base of the tentacles (arrows).

#### Thermal performance experimental design

Symbiotic and bleached polyps were moved to the experimental tanks on the day after the last menthol exposure and allowed to recover and acclimate for 10 days. The experimental system consisted of eight 5-L glass tanks (20 cm × 30 cm) equipped with a small pump (Resun SP-500) in a common temperature-controlled water bath (26 °C). White fluorescent lamps were used to maintain the light cycle and intensity consistent with long-term rearing conditions. Bleached and symbiotic polyps were kept in separate tanks, with four replicate tanks per treatment filled with FASW. Each tank contained one polyp from each colony (n = 3) that were used for the thermal performance experiment. Additional polyps were kept in each tank as back-up (Fig. S3). Polyps were fed each day after the end of the dark incubation, followed by 10 % water change after 2-3 h. Each polyp was fed one small frozen adult Artemia pipetted directly on top of the oral opening. The polyps were cleaned every two to three days after the incubations to remove fouling that would interfere with respirometry measurements.

To test the amenability to experimental handling of single polyps and to compare metabolic rates of bleached and symbiotic polyps, we conducted a thermal performance experiment. Photosynthesis and respiration of *G. fascicularis* polyps were measured across a 12 °C temperature range over 10 days. For this, the polyps were acclimated to a different temperature each day, in the following order: 20, 22, 23, 24, 25, 26, 27, 28, 30 to 32 °C. The temperature of the water bath was changed overnight, and incubations took place the subsequent day at the respective temperature.

#### Photosynthesis and respiration of symbiotic and bleached polyps

We measured oxygen evolution of individual *G. fascicularis* polyps in light and dark incubations to calculate net photosynthesis (PN) and dark respiration (R) respectively. For these measurements four bleached and four symbiotic polyps from three colonies were used (n = 12). Every day, each polyp was placed in individual 160 mL glass incubation jars (Weck, Germany), equipped with a magnetic stirrer (~5 cm s^−1^). Three additional jars containing only seawater and stirring bars were used to control background biological activity. Light intensity of 230 - 280 μmol photons m^−2^ s^−1^ during light incubations (ATI 80W Aquablue Special) and temperatures were maintained through thermostat-controlled water baths placed on a custom-made multiplate stirring system (Rades et al. 2022).

Dissolved oxygen (DO) was measured at the start and at the end of each incubation using a handheld multiparameter probe (WTW Multi 3620 IDS set). Light incubations lasted 2 h around mid-day followed by 1.5 h of dark incubation. Net photosynthesis and respiration were calculated using the following formula (Schneider and Erez 2006):

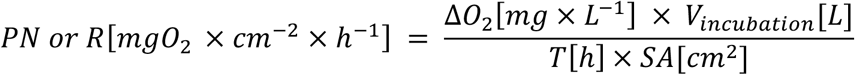

Where ΔO_2_ is the difference in dissolved oxygen between the end and the start of the incubation (DO_end_ - DO_start_) corrected for the controls’ ΔO_2_. This is multiplied by the volume of the incubation chambers in L (V_incubation_), and divided by incubation time (T) in hours and polyp surface area (SA) in cm^2^.

#### Surface area measurements

Polyp surface area was measured through photogrammetry. Between 40 to 50 pictures of each polyp were taken with a phone camera two days before the beginning of the incubations. These were used to create 3D models with 3DF Zephyr Free (v.4.523), cleaned on Artec3D (Studio 11 Professional v.11.2.2.16), and loaded on MeshLab (v.2016.12, Cignoni et al. 2008) for size scaling and calculation of live coral surface area.

#### Thermal performance data analysis

All analyses were conducted in the R statistical environment (v.4.1.0, R Core Team 2021) and plotted with ggplot2 (v.3.3.5, Wickham 2016). The effects of symbiotic state (‘symbiotic’ vs. ‘bleached’) or colony identity on net photosynthesis (PN) and respiration (R) were analyzed using linear mixed-effect models considering light and dark incubations separately. For this, ‘symbiotic state’, ‘colony identity’, and ‘temperature’ were set as fixed factors, and ‘polyp identity’ as random factor. The package ‘lmerTest’ (v.3.1-3, Kuznetsova et al. 2017) was used to construct the models and calculate p-values. The package ‘performance’ was used to check the residuals and to compare alternative models (v.0.7.3, Lüdecke et al. 2021). Differences between colonies were tested as pairwise comparisons of estimated marginal means (Searle et al. 1980) with Bonferroni correction with the ‘emmeans’ package (v.1.6.2-1).

### Symbiosis reestablishment after menthol bleaching

#### Native Symbiodiniaceae characterization

*G. fascicularis* colonies were collected in March 2019 from Crescent Bay in Hong Kong. DNA was successfully extracted from two colonies using the Qiagen DNeasy 96 Blood & Tissue kit. The ITS2 marker region was amplified with the primers SYM_VAR_5.8S2 and SYM_VAR following the PCR protocol by Hume et al. (2018), followed by paired-end sequencing on the Illumina MiSeq platform (2 × 300 bp). Raw sequencing data was analyzed using the SymPortal workflow remote instance (Hume et al. 2019), with results reported as post-MED ITS2 sequences in relative abundances.

#### Menthol bleaching and inoculation of cultured Symbiodiniaceae

We explored the possibility of returning adult corals to the symbiotic state after menthol bleaching. For this, we inoculated menthol-bleached coral fragments from Hong Kong with cultured Symbiodiniaceae. Bleaching followed the same protocol described for the Red Sea colonies. Two species of symbionts, *Cladocopium goreaui* and *Durusdinium trenchii* were chosen for targeted exposure of the bleached fragments, as species from both of these genera are known to form stable symbiosis with *G. fascicularis* across the South China Sea (Tong et al. 2017). Batch cultures of *C. goreaui* (AIMS-SCF-055, ITS2 type-sequence C1, isolated from *Acropora tenuis*) and *D. trenchii* (AIMS-SCF-088, ITS2 type-sequence D1-4, isolated from *A. muricata*) were obtained from the Symbiont Culture Facility at the Australian Institute of Marine Science and maintained in f/2 media (Guillard 1975) under the same conditions as the corals. A total of 16 fragments were inoculated with a concentration of 200 symbiont cells ml^−1^ in each 6-L aquarium. The concentration of symbionts was divided in a 90:10 ratio (180 cells ml^−1^ *D. trenchii* : 20 cells ml^−1^ *C. goreaui*). This ratio was chosen because members of *Cladocopium* are the most prevalent symbionts found in *G. fascicularis* in Hong Kong (Huang et al. 2011; Tong et al. 2017) and across its wide geographic range from the Red Sea to the South China Sea, where *Durusdinium* is only occasionally found and only in comparably warm habitats (Dong et al. 2009; Ziegler et al. 2017; Zhou et al. 2017). The symbiont aliquots were centrifuged to remove the culture media and washed once with FASW before final resuspension in 10 mL FASW. The bleached fragments were first fed before being exposed to the chosen symbionts. For this, a suspension of ReefRoids in ASW was pipetted directly into the oral opening of each polyp. The same process was then immediately repeated with the suspended symbionts. Aquarium filters were turned off for 1 h to allow the corals to feed and to prevent removal of symbiont cells from the water column. Fragments were inoculated twice a week for 6 weeks, until symbiosis reestablishment was visually confirmed. Symbiodiniaceae populations *in hospite* were then characterized with fluorescent in-situ hybridization (FISH) and flow cytometry (McIlroy et al. 2020).

#### Fluorescent In-Situ Hybridisation (FISH)

After 6 weeks of inoculation, a single polyp was removed from each fragment with a chisel. Tissue was removed from the skeleton with an airbrush containing deionized water. The resulting slurry was homogenized using a 20-μl pipette tip. 1.5 mL of homogenate was transferred to a 2-mL Eppendorf tube and centrifuged at 500 rpm for seven minutes, after which the supernatant was discarded. The remaining pellet was washed once before being preserved in 5X SET buffer and stored at −80 °C for further analysis.

Samples were prepared following an adapted protocol from McIlroy et al. (2020). Briefly, after washing with 5X SET in IGEPAL at 0.4 and 0.1 % vol/vol samples were split equally between three 1.5-mL black Eppendorf tubes. Each sample had one tube treated with a SymC probe, one tube with a SymD probe and one without any probe as a control. Both SymC (5’-CTACCCAAGAACTTGCAGG-3’) and SymD (5’-CCACCCAAGAACTCGCGTG-3’) probes were designed to have genus-level specificity for *Cladocopium* and *Durusdinium*, respectively, and were labelled at 5’ end with Alexa Fluor ® 532. After overnight hybridization at 45 °C (5X SET, 0.1 % vol/vol IGE-PAL, 10 % vol/vol formamide and 100 pmol of the relevant probe), samples were washed once in warm 1X SET, vortexed and resuspended in 500 μl of 1X SET for same-day cytometric analysis on a BD FACSAria Fusion flow cytometer (BD Biosciences, CA). A gating hierarchy was used to isolate cells of interest. Cells were screened for size and presence of doublets (multiple cells stuck together) based on light-scatter parameters (forward-scatter (FSC) and side-scatter (SSC) (Fig. S6a-c). Probe-positive cells were then distinguished based on their position within the 582m vs SSC-A scatterplot (Fig. S6d,e). FlowJo™ Software (v.10.8.1, BD Life Sciences) was used to determine the percentage of probe-positive cells (either *Cladocopium* or *Durusdinium*), to identify the presence and the relative abundance of each algal genus within the sample.

### Completing gametogenic cycles and *ex-situ* spawning

*Galaxea fascicularis* in the GBR spawns between October and December, 0.05-200 minutes after sunset (MAS) and 1 – 8 nights after full moon (NAFM) (Babcock et al. 1986; Baird et al. 2021). To determine if the *ex situ* reproductive pattern remained in synchrony with the wild we followed our colonies over three consecutive reproductive seasons (2019 – 2021).

To ascertain gamete development in each colony, two months prior to predicted spawning, and two to four days prior to the full moon, whole polyps were removed from each colony and longitudinal sections imaged (Canon 5d MKIII, Fig. 5a,d). Based on the stage of gamete development in these sections *ex situ* spawning activity was predicted.

To record spawning observations each year, colonies were observed with red head torches during the predicted spawning months. Observations commenced two NAFM and 30 mins prior to the predicted spawning time, during which time the broodstock aquariums were isolated from the filtration system. In addition, all internal water pumps were also turned off leaving the aquariums’ water static. Observations continued over multiple consecutive nights until all colonies that had developed gametes, during that reproductive season, spawned. Spawning date and time of each colony was recorded, with onset of spawning being denoted as the time of first gamete release. Following release, gametes were pooled to allow fertilization to occur, and subsequent embryos were reared and larvae settled following the method described by Craggs et al. (2019).

## Results

### Production and long-term rearing of bleached polyps

We produced single polyps (clonal replicates) of *G. fascicularis* colonies from the Red Sea and bleached them with menthol. All polyps (n = 24, including additional polyps not used in this experiment) survived the menthol treatment and appeared visually completely white. Similarly, all menthol-bleached coral fragments from Hong Kong (n = 16) survived the treatment. Visual inspection under a fluorescent stereomicroscope revealed no detectable algal cells in the bleached polyps (Red Sea colonies, Fig. 2a), but inspection at higher magnification (compound fluorescent microscope, ×400) revealed the presence of few scattered algal cells, limited to the base of the tentacles (Fig. 2b).

All polyps that were employed in the thermal performance experiment (n = 12) remained visually completely bleached and viable throughout the 10 days of temperature treatment, with the exception of two polyps that appeared dead on the 7^th^ and 8^th^ day of incubation, respectively. These polyps showed atrophic and unresponsive tentacles, and overall thinning of the soft tissue until it was not recognizable. Overall, we successfully maintained the bleached polyps for three weeks from the termination of the menthol treatment.

### Amenability to experimental manipulation and broad thermal tolerance

Symbiotic (untreated) polyps had positive net photosynthesis across all temperatures from 21 to 32 °C (Fig. 3). Net photosynthesis peaked at 23 °C (mean 13.05 μg O_2_ cm^−2^ h^−1^) and reached the minimum at 32 °C (mean 3.26 μg O_2_ cm^−2^ h^−1^). In contrast, respiration overall increased with temperature with minimum values at 21 °C (mean 5.77 μg O_2_ cm^−2^ h^−1^) and maximum at 32 °C (mean 25.70 μg O_2_ cm^−2^ h^−1^; Fig. 3). Bleached polyps, as expected by their lack of photosymbionts, exhibited a net oxygen depletion. Similar to symbiotic polyps, respiration in bleached polyps increased with temperature, both in light and dark incubations (Fig. 3). Overall, respiration in symbiotic polyps was higher than in bleached polyps (P_lmer_ < 0.0001).

**Fig. 3.**
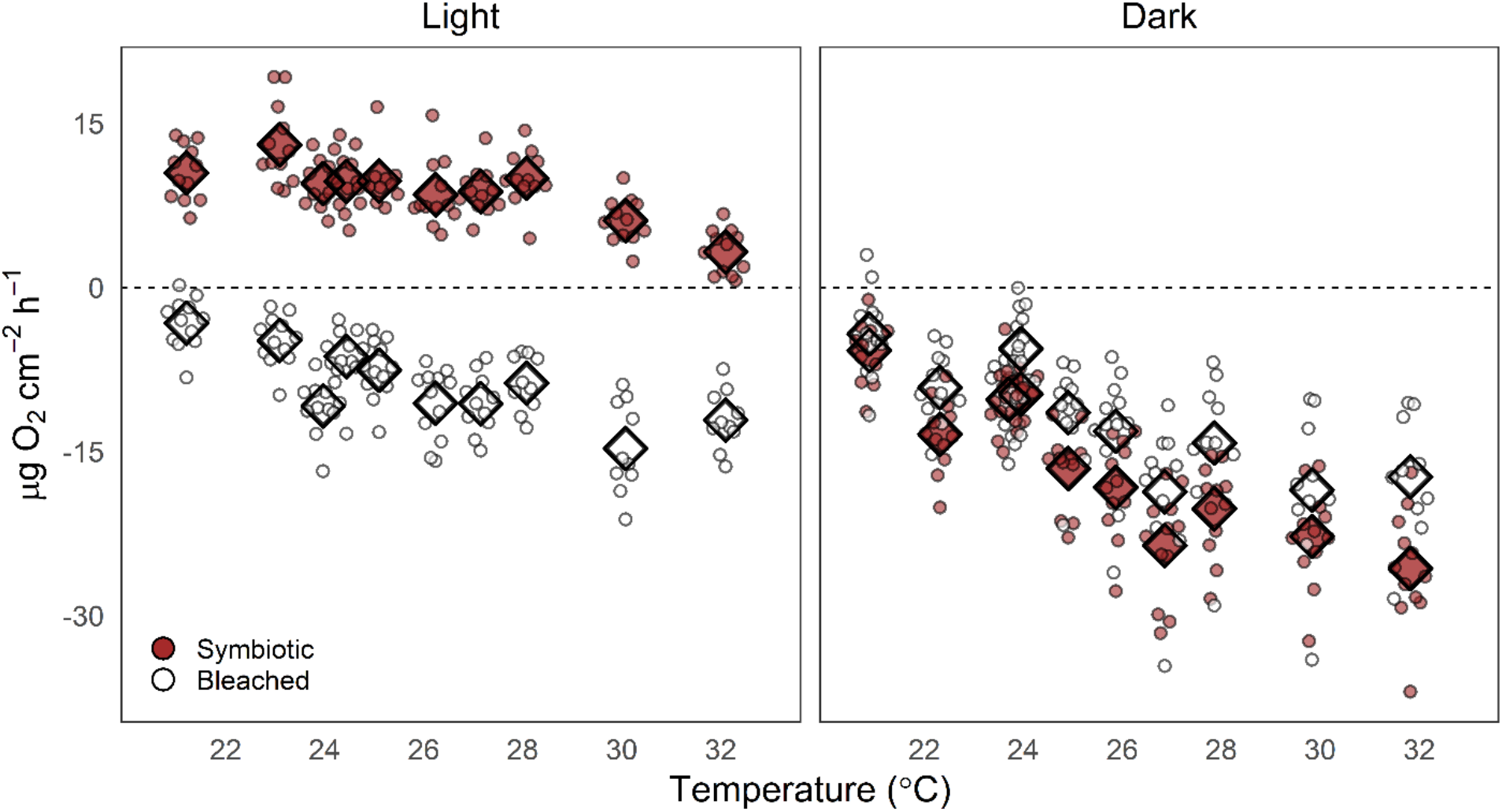
Net photosynthesis and respiration rate of symbiotic states in *Galaxea fascicularis*. Symbiotic (brown symbols) and bleached (white symbols) polyps in light and dark incubations. Noise added (jittering) for ease of visualization, means per group depicted as diamonds. n = 12

While net photosynthesis was similar between colonies (P_lmer_ > 0.05), respiration was significantly higher in one colony (P_lmer_ < 0.001) compared to the other two (RS2-RS1: P_Bonf_ < 0.01; RS2-RS3: P_Bonf_ < 0.01; RS1-RS3: P_Bonf_ > 0.05; Fig. S4).

### Symbiosis re-establishment after bleaching

Inoculation with cultured symbionts resulted in successful symbiosis re-establishment for 100 % of the samples (n = 16), which were macroscopically visibly symbiotic. Of these, nine samples were successfully FISH hybridized, with the genus-specific probes identifying the presence of both target genera in each sample. Individual variation in the relative ratio of each genus was evident among the fragments, with *Durusdinium* contributing to 4 – 63 % of cells and *Cladocopium* to 37 – 82 % (Fig. 4, Tab. S2). A small, but significant proportion of symbionts remained unlabeled in some samples. These may represent additional symbiont genera or be the result of variable probe efficiency. *G. fascicularis* colonies from the same location were found to harbor *Cladocopium* symbionts only (dominant ITS2 sequence types: C1 and C1c, Tab. S3).

**Fig. 4.**
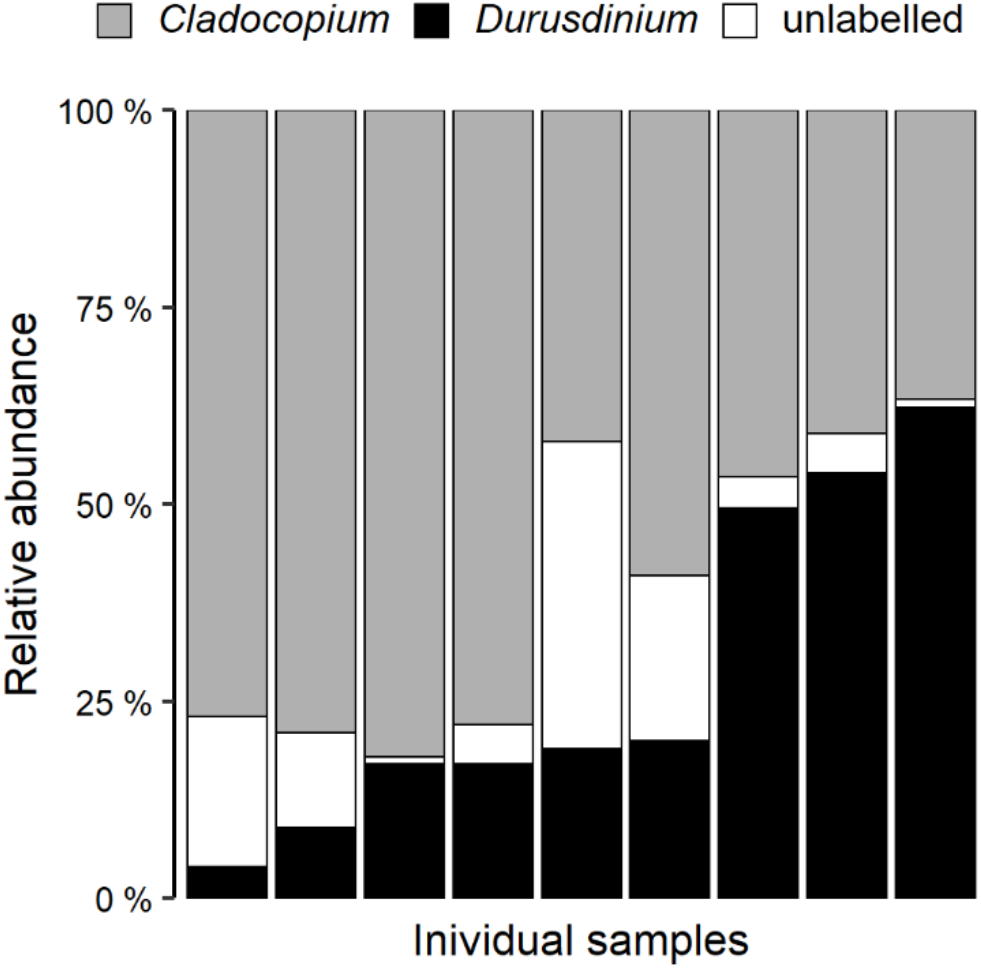
Symbiosis reestablishment in menthol-bleached adult corals inoculated with cultured Symbiodiniaceae of the genera *Cladocopium* and *Durusdinium*. The presence and relative abundance (%) of both genera was confirmed and quantified through fluorescent in-situ hybridization (FISH) and flow cytometry.

### Ex-situ spawning

Spawning was monitored each year over the three-year observation period for all 35 colonies. Annual fecundity, as a percentage of colonies completing a full gametogenic cycle, was on average 56.2 % during these three spawning cycles (2019: 48.6 %, 2020: 45.7 %, and 2021: 74.3 %).

Spawning occurred in synchrony with wild predictions, with gametes being released between October and January each year, between 5 - 259 MAS and 4 - 13 NAFM (Fig. 5b-c & e-f, Fig. 6). Following embryo rearing, Symbiodiniaceae were observed within primary polyps 14 days post settlement (Fig. 5g). These originated from the adult colonies housed in the same system and taken up via the water column. Polyps were fully developed 40 days post settlement (Fig. 5h) and grew into small colonies within the first six months (Fig. 5i).

**Fig. 5.**
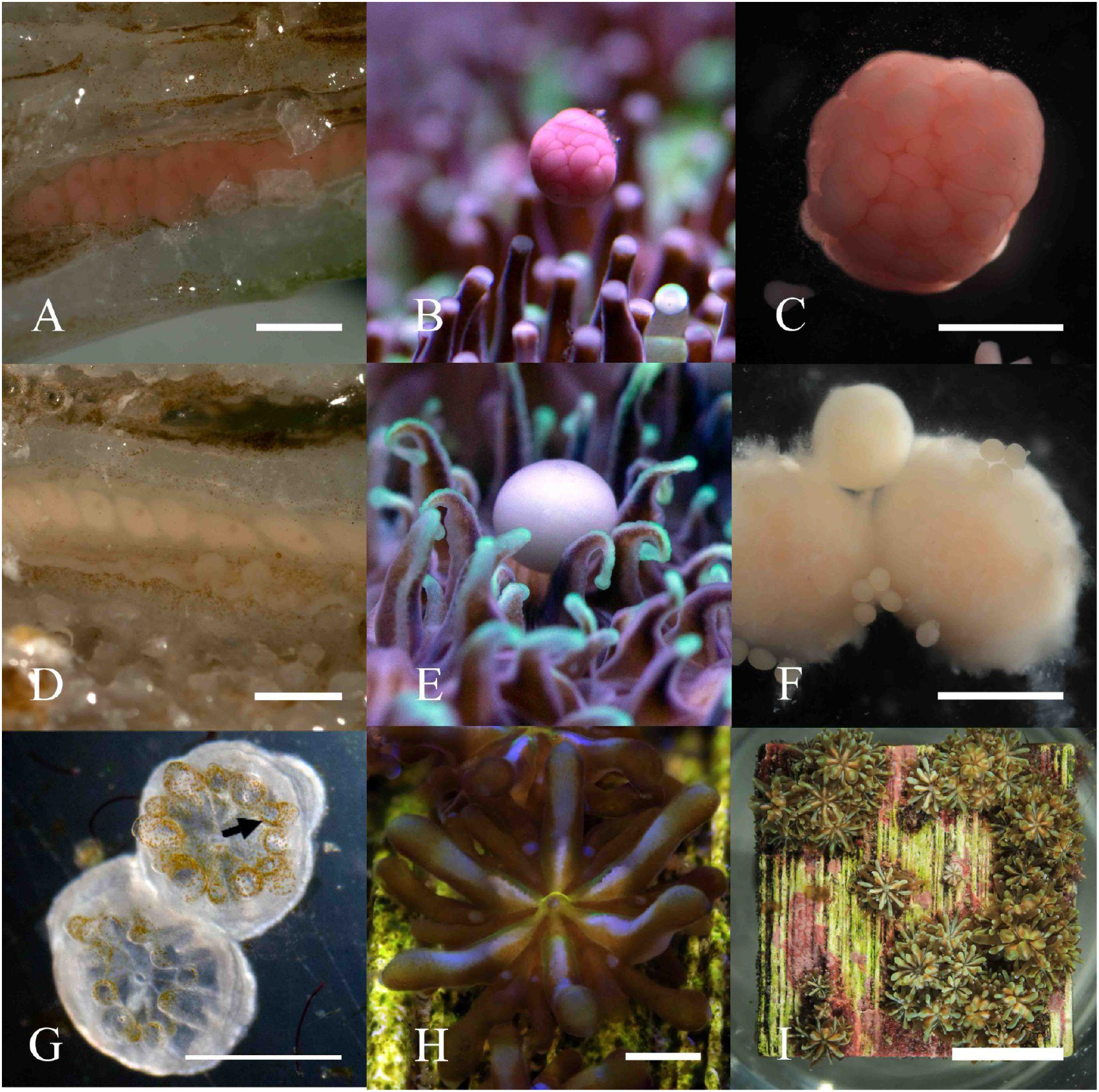
Representative images of female and male colonies of *Galaxea fascicularis* spawning *ex situ* and gamete development. **a** Longitudinal section of female polyp with pinkish-red pigmented oocytes. **b** Female colony releasing oocyte bundle *ex situ*. **c** Close-up of female bundle post release. **d** Longitudinal section of male polyp with white oocytes. **e** Male colony releasing oocytes/sperm bundle *ex situ*. **f** Close-up of male gamete bundles post release undergoing disassociation. **g** Newly settled *G. fascicularis* primary polyps showing initial Symbiodiniaceae (arrow) uptake 14 days post settlement. **h** Primary polyp 40 days post settlement. **i** Multiple colonies six months post settlement. Scale bar: a-h = 1 mm, i = 1 cm.

**Fig. 6.**
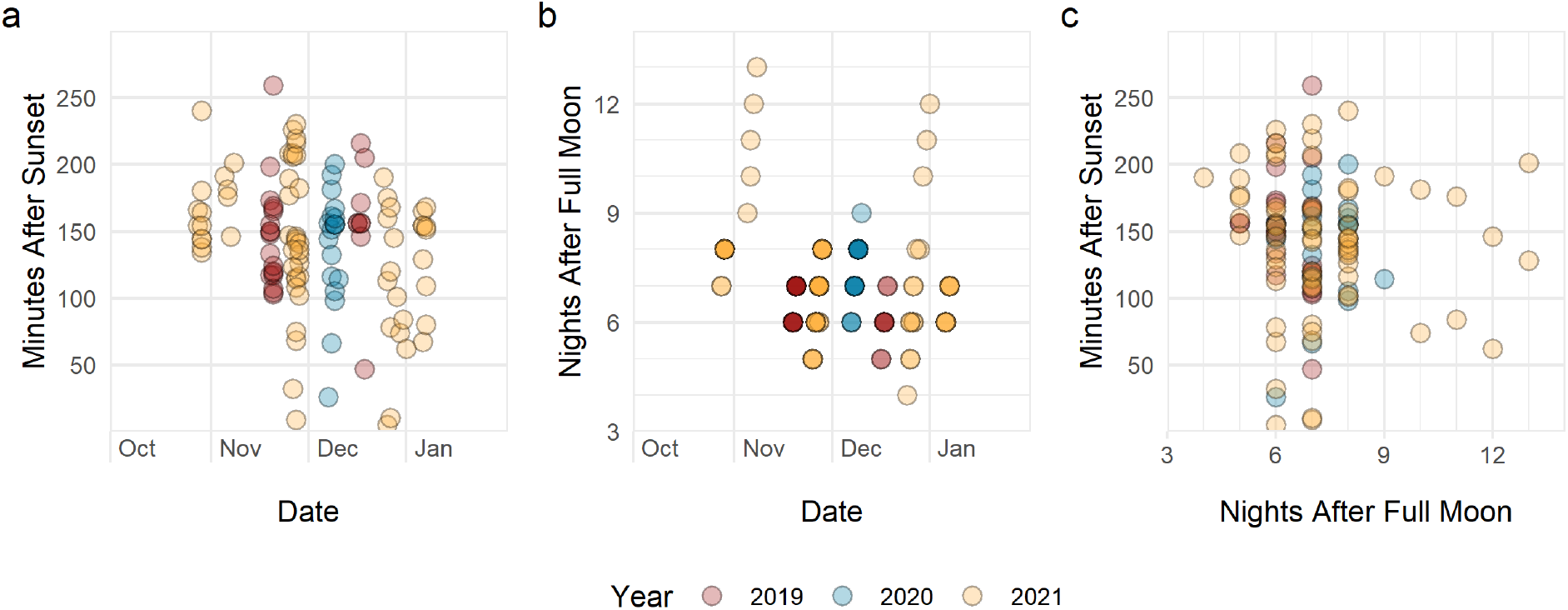
*Galaxea fascicularis ex situ* spawning periodicity over three spawning cycles (2019 – 2021). **a** Spawning time, recorded as the first gamete release in Minutes After Sunset (MAS) by date. **b** Night After Full Moon (NAFM) of gamete release by date. **c** Comparison of MAS and NAFM values over three *ex situ* spawning cycles. 2019 n = 28, 2020 n = 19, 2021 n = 69.

## Discussion

Here, we explored the potential of *G. fascicularis* as a novel coral model for symbiosis research by assessing its amenability to aquarium rearing, experimental handling, and symbiosis manipulation.

### Successful bleaching with menthol

We showed that *G*. *fascicularis* can be readily and reliably rendered aposymbiotic through menthol bleaching, similar to other cnidarian model systems (e.g., Matthews et al. 2015; Röthig et al. 2021). We lowered the menthol concentration in the protocol developed by Wang et al. (2012) to further limit stress on the host, and indeed all coral polyps survived this bleaching procedure on two independently replicated occasions. Overall, menthol treatment was very effective, with only a few algal cells remaining at the base of the tentacles. From a photophysiological perspective, these scant remnants can be considered negligible, as illustrated by the light and dark oxygen production data. Furthermore, bleached polyps remained aposymbiotic for weeks and we therefore considered this bleaching protocol suitable for experimental investigation of aposymbiotic *G*. *fascicularis* in physiological studies and symbiosis manipulation.

Notably, besides Symbiodiniaceae, reef-building corals also associate with other microorganisms (i.e., bacteria, viruses, and other unicellular algae and protists) which constitute the so-called coral holobiont (Rosenberg et al. 2007). Although this has only been appreciated more recently, holobiont composition and dynamics are now understood to underlie coral health and adaptability (Pogoreutz et al. 2020). As menthol has photo-inhibitory (Wang et al. 2012; Clowez et al. 2021) and antimicrobial properties (İşcan et al. 2002), further studies should investigate the effects of menthol bleaching on the remaining members of the microbiome.

### Tractability of symbiotic and aposymbiotic *G. fascicularis*

Tractability, as amenability to experimental work and laboratory conditions, is one of the most important requirements for model organisms. The *G*. *fascicularis* polyps and fragments could be maintained for three and ten weeks respectively in simple systems consisting of small independent tanks (5-6 L) with regular feeding and basic water quality care. These time frames match the requirements for most experimental designs, and we believe could be further extended. Indeed, symbiotic individuals looked healthy and responsive throughout the whole period, testifying to a comparatively hardy coral with only low demands on rearing conditions. Additionally, the positive net photosynthetic rate across a large temperature range confirmed the broad thermal tolerance of this species (Al-Horani 2005; McIlroy et al. 2019), which was retained in the simplified single polyp application.

Moreover, bleached polyps could be maintained in the simple experimental system for three weeks even with the additional stress of the thermal experiment. However, they eventually showed signs of deteriorating condition indicating that not all nutritional needs were met. Thus one priority is to develop bespoke feed for long-term rearing of bleached corals (Wang et al. 2012).

### Thermal performance of symbiotic and aposymbiotic *G. fascicularis*

Studies that compare symbiotic and aposymbiotic coral holobionts are particularly informative for understanding coral response to climate change, and therefore we investigated the comparability between symbiotic and aposymbiotic *G*. *fascicularis* polyps. As expected, these differed in their photosynthetic rates, where the latter were net oxygen consumers even under illumination. Overall, symbiotic polyps had higher respiration rates than aposymbiotic polyps, which could be ascribed to the additional metabolic burden of the associated Symbiodiniaceae, as well as a greater metabolic capacity derived from the availability of photosynthates (Muscatine et al. 1981; Al-Horani et al. 2003). For all polyps, respiration rates overall increased with temperature in line with optimum thermal performance of corals and macroalgae from the central Red Sea (27-33 °C) (Anton et al. 2020).

Interestingly, we observed intraspecific variability in physiological performance. One colony (RS2) had significantly higher respiration rates than the others (Fig. S4). This manifested exclusively under dark conditions and was therefore more likely linked to host rather than symbiont identity. This variability warrants further investigations to better characterize host genotypes, as in the long term it will be necessary to establish representative host clonal lines.

### Symbiosis reestablishment in adult corals

We showed that adult bleached *G. fascicularis* can successfully reestablished symbiosis with cultured Symbiodiniaceae. The experimental repopulation of adult corals with cultured non-native symbionts has only recently been demonstrated (Scharfenstein et al. 2022), and it is considered particularly relevant in the context of symbiosis manipulations to enhance heat tolerance in reef-building corals (van Oppen et al. 2015). Notably, here this was further explored with two non-native algal cultures during simultaneous exposure. The first, *D. trenchii* (ITS2 type D1-4), has never been found in *G. fascicularis* from Hong Kong (Huang et al. 2011; Tong et al. 2017) nor in our characterization of Symbiodiniaceae from that site specifically (two other colonies from the same location, Tab. S3) and can therefore confidently be considered to be non-native. The second, *C*. *goreaui* (ITS2 type C1), was isolated from the coral species *A. tenuis* from Australia. Even though the dominant C1 ITS2-sequence is also present in *G. fascicularis* from Hong Kong, the high coral host-Symbiodiniaceae specificity, large radiation of C1-like Symbiodiniaceae (LaJeunesse 2005; Thornhill et al. 2014), and the consistent occurrence of the minor sequence C1c in addition to C1 in our samples, implicate the *C. goreaui* culture as non-native in *G. fascicularis*. However, the higher similarity with the native symbionts likely explains the more efficient establishment of the *C. goreaui* culture than the *D. trenchii* culture despite a reversed (10:90) ratio in supply.

*Galaxea fascicularis* has generally been reported to host taxa from the genera *Cladocopium* and *Durusdinium*, both exclusively and simultaneously, and with an equatorial-ward shift from *Cladocopium* dominance to *Durusdinium* dominance (Huang et al. 2011, and references therein). At Hong Kong’s latitude and at our rearing temperature, *G*. *fascicularis* is expected to be exclusively associated with *Cladocopium* (Huang et al. 2011; Tong et al. 2017; Zhou et al. 2017), yet the targeted inoculation produced mixed-genus associations. This demonstrated how targeted inoculation can successfully shift Symbiodiniaceae community composition towards a higher proportion of heat tolerant strains without thermal treatment (Silverstein et al. 2015). Although the longevity of these partnerships is still unclear, these are promising results for future development of *in hospite* studies on the role of temperature on inter-partner symbiosis dynamics and symbiont recognition mechanisms (Bove et al. 2022). The success and potential insights from such experimental approaches will rely on future efforts to establish a variety of coral host-specific Symbiodiniaceae cultures (including the establishment of specific cultures from *G. fascicularis*) for further study.

Of note, repopulation is possible even with lower concentrations of Symbiodiniaceae, as we observed from earlier qualitative trials (details in Supplementary Text). There, we exposed the menthol bleached polyps after they were used in the thermal performance experiment to Symbiodiniaceae through the water medium (symbionts shed by nearby symbiotic colonies) or additionally in combination with a one-time inoculum of freshly isolated symbionts from symbiotic conspecifics. The success rates of these qualitative trials of ~30 % and 43 %, respectively, were lower than in the Hong Kong experiment (100 %) yet remarkable considering the poorer health conditions of the polyps at the time of re-exposure to symbionts (i.e., after repeated incubations through the 21-32 °C temperature excursion), and the expectedly smaller number of symbionts available for uptake. In these trials, bleached hosts retained the ability to feed, which is known to facilitate symbiont uptake (Fitt and Trench 1983) and likely contributed to symbiosis re-establishment. In the long-term, the polyps that reestablished symbiosis grew into small colonies and became visually indistinguishable from the untreated controls (Fig. S5). Taken together, these observations attest to the suitability of *G. fascicularis* to experimental symbiosis manipulation.

### *Ex situ* spawning

*Galaxea fascicularis* is a good model species candidate for sexual reproductive work due its high survivorship in aquarium conditions. As demonstrated herein, a combination of seasonal programming and husbandry enabled the species to complete a full gametogenic cycle *ex situ* over multiple years, which closely mimics that observed in the wild (Babcock et al. 1986; Baird et al. 2021). Furthermore, we showed *in vitro* fertilization, embryological development, and *ex situ* larval settlement resulting in spat of known ages. As the age at which the species reaches sexual maturity is currently unknown, future studies should focus on this aspect, which is also important for producing an F2 generation and for closing the species life cycle *ex situ* (Craggs et al. 2020). Access to a F2 generation of known parental lineage will provide a platform to conduct experiments in areas such as mapping quantitative trait loci (Zhang 2012) or phenotypic traits such as growth, disease resistance, or thermal tolerance (van Oppen et al. 2015). As *Galaxea* must acquire their symbionts from the seawater (horizontal transmission, with symbionts visible after the development of tentacles in settled polyps, Lin et al. (2022)), *ex situ* spawning also opens possibilities for studies on the dynamics of initial symbiosis establishment and how this is affected by symbiont identity, heritability in symbiont selection, and for rearing of axenic or gnotobiotic coral lineages, paralleling procedures established for *Hydra* (Fraune et al. 2015).

## Conclusions

The study of coral-algal symbiosis has been constrained by the difficulties of maintaining corals *ex situ* and of manipulating the coral-algal association to unravel symbiosis functioning. Here, we showed that adopting *G. fascicularis* as a model system can alleviate these limitations. Its demonstrated compatibility to rearing in simplified systems and experimental applications, together with the repeated spawning in aquaria, confirm the high tractability of this coral species. Further, the possibility to readily obtain aposymbiotic individuals suitable for detailed study and symbiosis re-establishment set the stage for exciting future developments in coral symbiosis research which is necessary to understand corals’ potential to cope with changing oceans.

Besides these strengths, our study also highlights areas worth future efforts on the path to developing the *G. fascicularis* model system further. These include: i) better characterization of the effect of menthol on the rest of the coral microbiome (e.g., bacteria); ii) development of custom feed for long-term maintenance of bleached individuals; iii) assessment of long-term persistence of non-native Symbiodiniaceae, and testing of a variety of symbiont strains to understand the breadth of symbiont diversity that can be accommodated by the *G. fascicularis* host; iv) further characterization of host colonies and genotypes with the aim of establishing clonal lines, and from these, isolate and establish Symbiodiniaceae cultures, mirroring for example the development of the Aiptasia model; and v) development of collaborative platforms and open-access community resources around this emerging model system to accelerate research and discovery.

## Acknowledgements

We thank Dr. Jafargholi Imani and Dr. Kathrin Ehlers (both JLU) for access and guidance in the fluorescence microscope facilities, André Dietzmann and Catarina P. P. Martins (both JLU) for assistance during experiments, and the Li Ka Shing Faculty of Medicine Core Facility (HKU) for the flow cytometry equipment.

This study is part of the ‘Ocean2100’ global change simulation project of the Colombian-German Center of Excellence in Marine Sciences (CEMarin) funded by the German Academic Exchange Service. We acknowledge financial support by the German Research Foundation (DFG, Project: 469364832) to MZ and Hong Kong Research Grants Council #17117221 to SEM and #17108620 to DMB.

## Data and code availability

Data and R scripts used for this study are available at: https://github.com/sPuntinG/Galaxea_Coral_Model/

## Author contributions

GP and MZ conceived and designed the study. GP, JC, RH, KEE, SM produced and analyzed data. MS, DMB, MZ contributed reagents/materials/analysis tools. GP and MZ wrote the first draft with contributions from all authors.

## Conflict of Interest

The authors declare no conflict of interest for this submission.

## Notes

### Competing Interest Statement

The authors have declared no competing interest.

### Summary of Updates

New Figure 1 added to summarize study design New Figure 6 added to visualize spawning periodicity Trials on qualitative symbiosis re-establishment moved to supplement Additional supplemental tables added to provide details on symbiosis reestablishment, new ITS2 sequencing of Symbiodiniaceae, and rearing conditions across labs

