## Supplementary material for "The reef-building coral *Galaxea fascicularis*: a new model system for coral symbiosis research"

#### Supplementary Figures

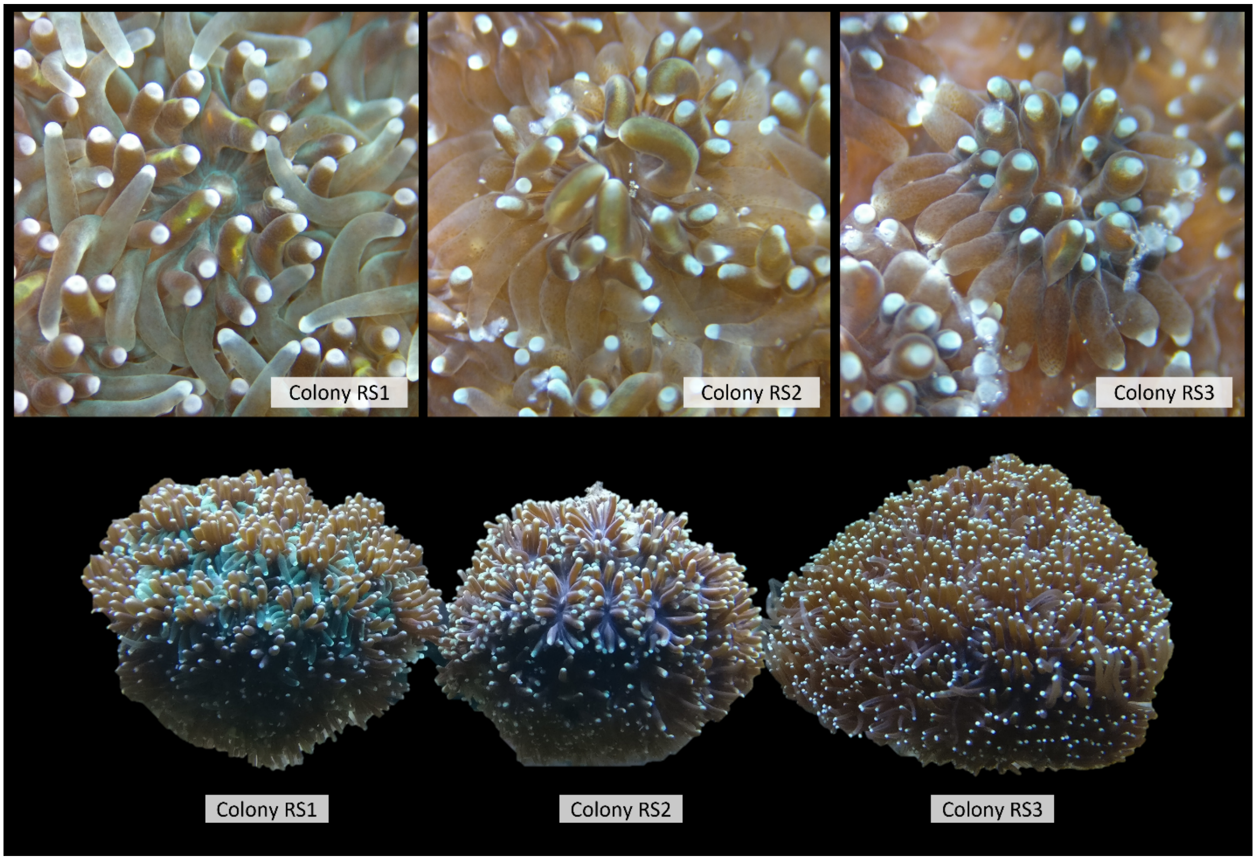


**Fig. S1 Three colonies of *Galaxea fascicularis* from the Red Sea** used in the thermal performance curve experiment showing slightly distinct coloration. Top: stereomicroscope with ×12.8 magnification (Leica, EZ4). Bottom: macroscopic pictures of the colonies in the aquarium tank, *Ocean2100* aquarium facility, JLU Giessen (Germany).

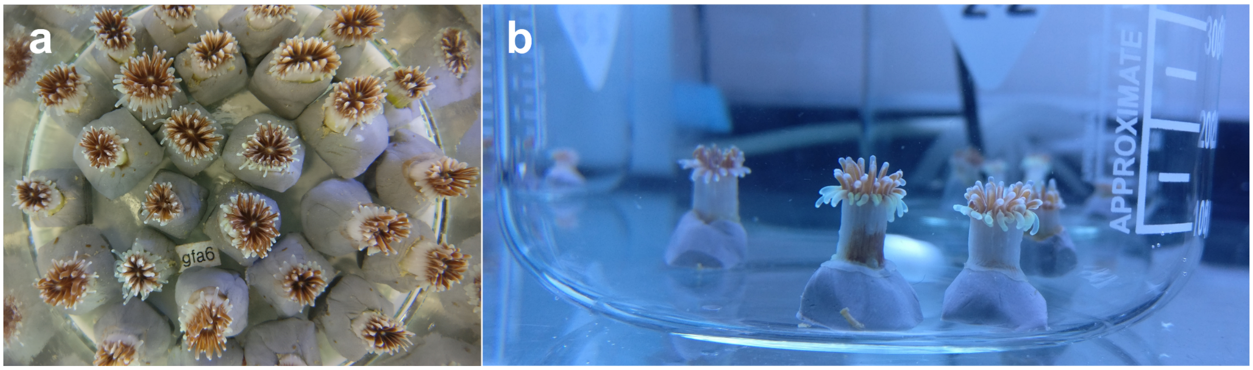


**Fig. S2 Examples of single polyps of *Galaxea fascicularis*.** **a** subsample of the polyps used for the thermal performance curve experiment; **b** polyps during menthol treatment in 1-L beaker with stirring.

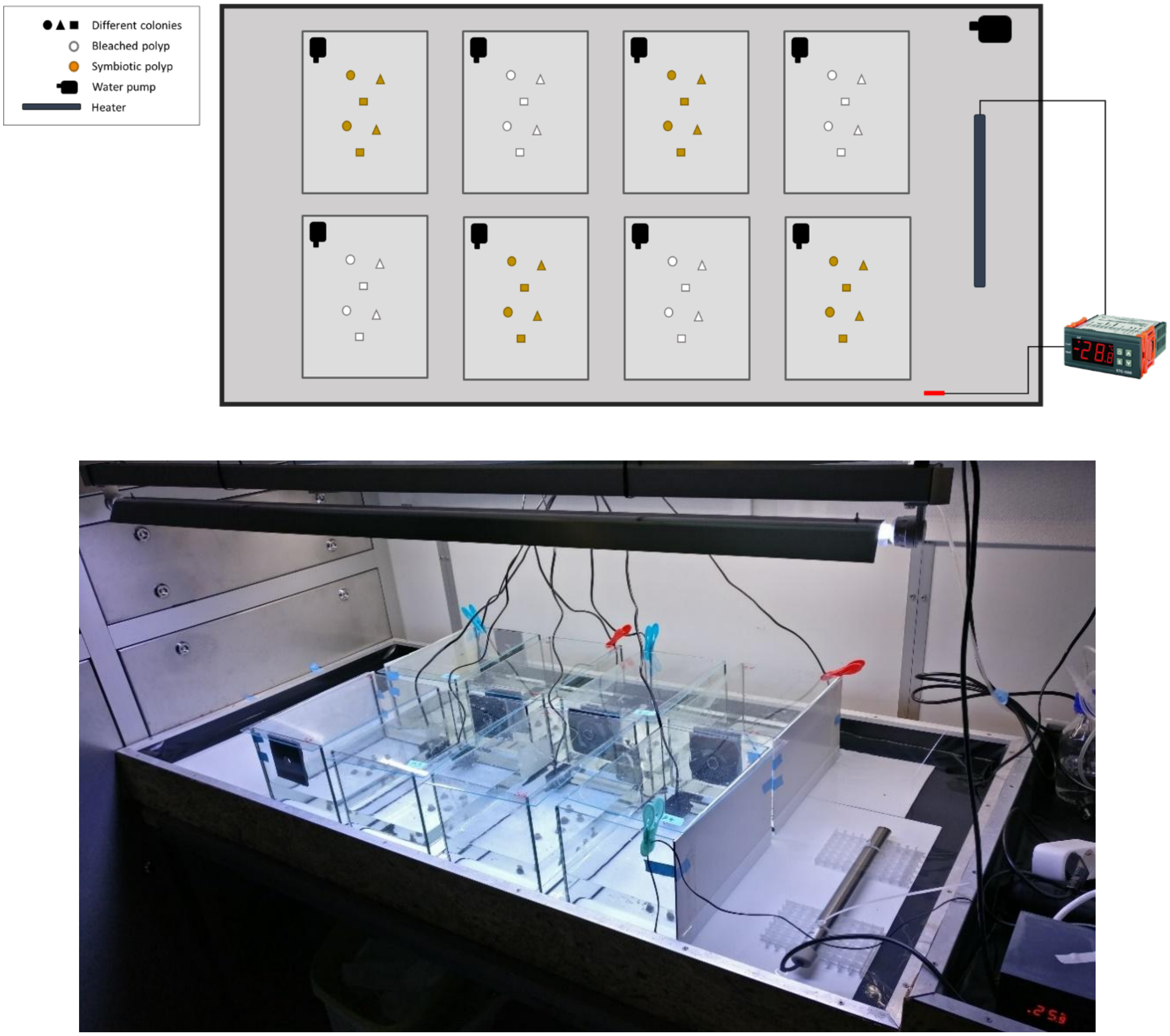


**Fig. S3 Simplified experimental set-up**. Schematic representation (top) and picture (bottom) of the experimental set up of the thermal performance experiment.

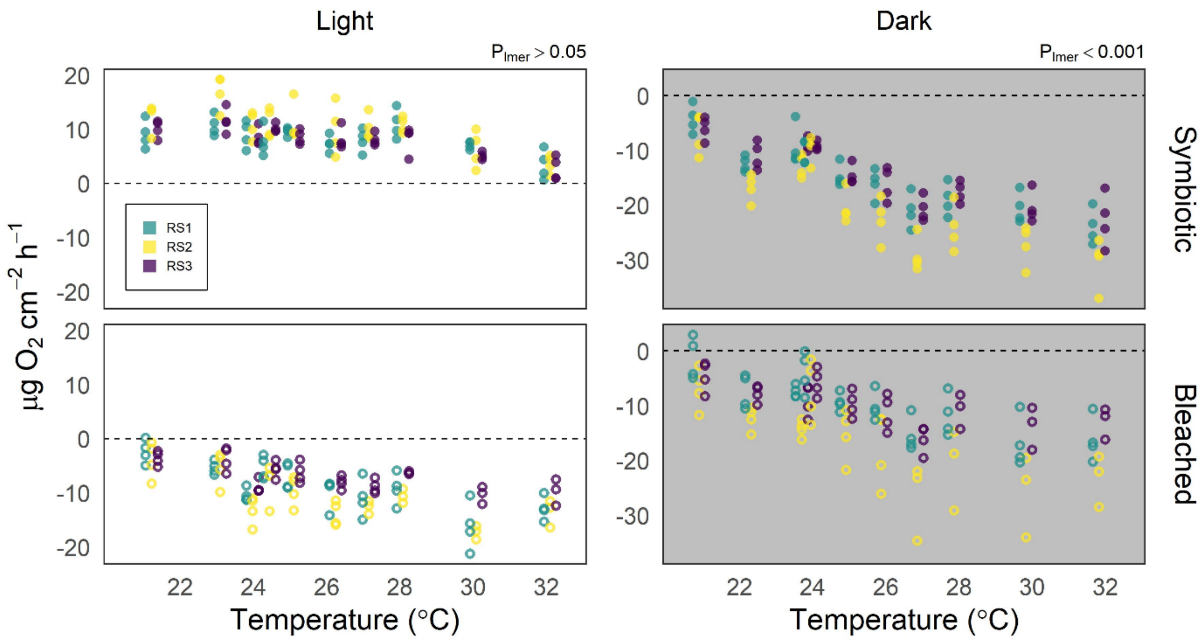


**Fig. S4 Net photosynthesis and respiration rates of symbiotic and bleached polyps of *Galaxea fascicularis* from three different colonies, in light and dark incubations**. Color indicates colony identity. Full circles: symbiotic polyps; hollow circles: bleached polyps. For each temperature step, points are dodged (by colony identity) for ease of visualization

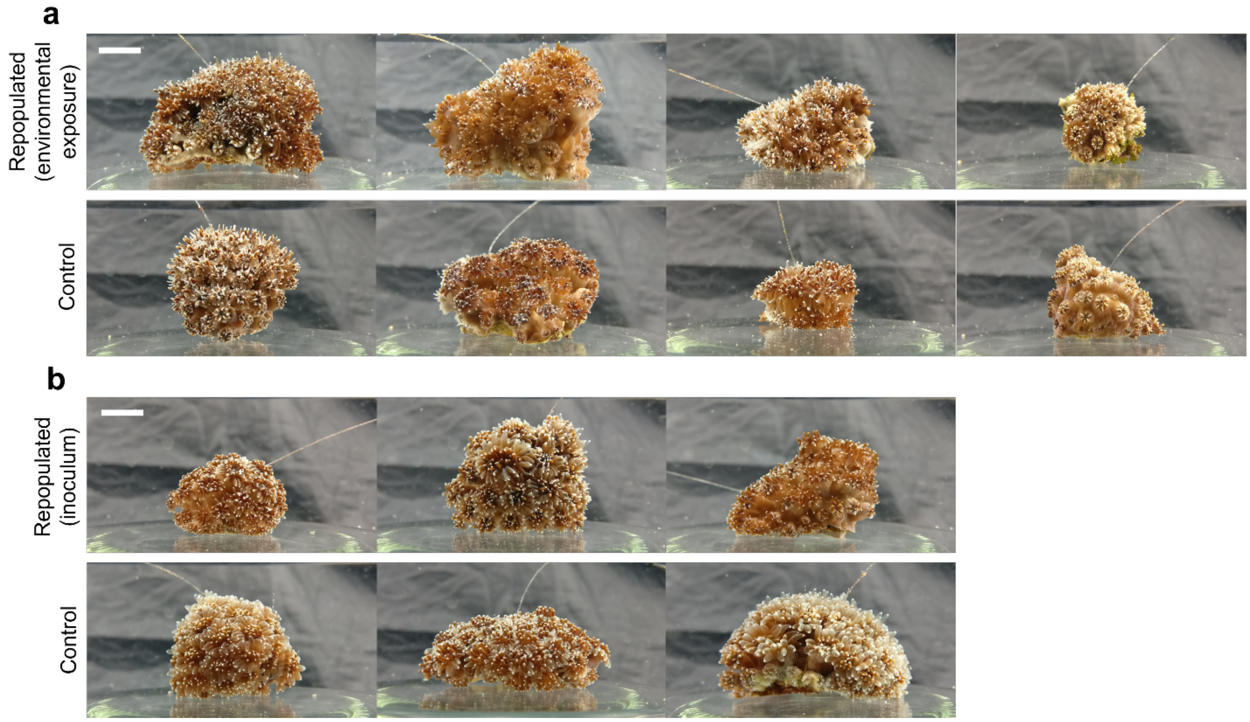


**Fig. S5 Small colonies of *G. fascicularis* originated each from a single menthol-bleached polyp following symbiosis reestablishment with Symbiodiniaceae**. **a** 'environmental' uptake and repopulation from Symbiodiniaceae acquired through the water column; **b** Symbiosis reestablishment in bleached polyps inoculated once with freshly isolated Symbiodiniaceae. Colonies grown from unbleached polyps shown as controls for each morphotype. Scale bar = 1 cm.

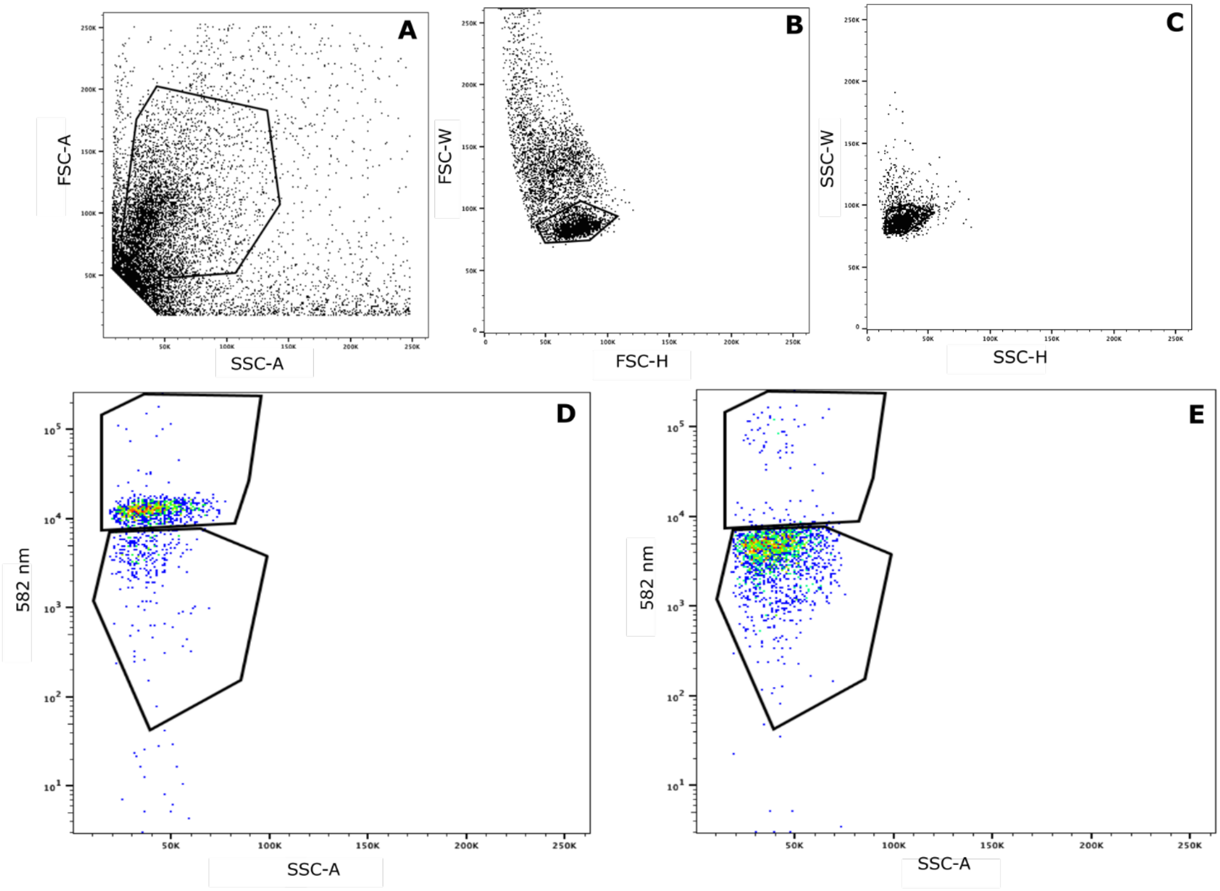


**Figure S6 Gating hierarchy for flow cytometric sorting of mixed symbiont communities.** **a** Cells of interest are isolated based on size (forward scatter; FSC-A) and shape (side scatter, SSC-A). Doublets, where multiple cells are analyzed in a single droplet, are removed by (**b**) secondary screening of forward scatter (width vs height; FSC-W vs FSC-H) and (**c**) side scatter (width vs height; SSC-W vs SSC-H). SymC- (**d**) or SymD-labelled (**e**) cells were then distinguished by increased fluorescence at 582nm.

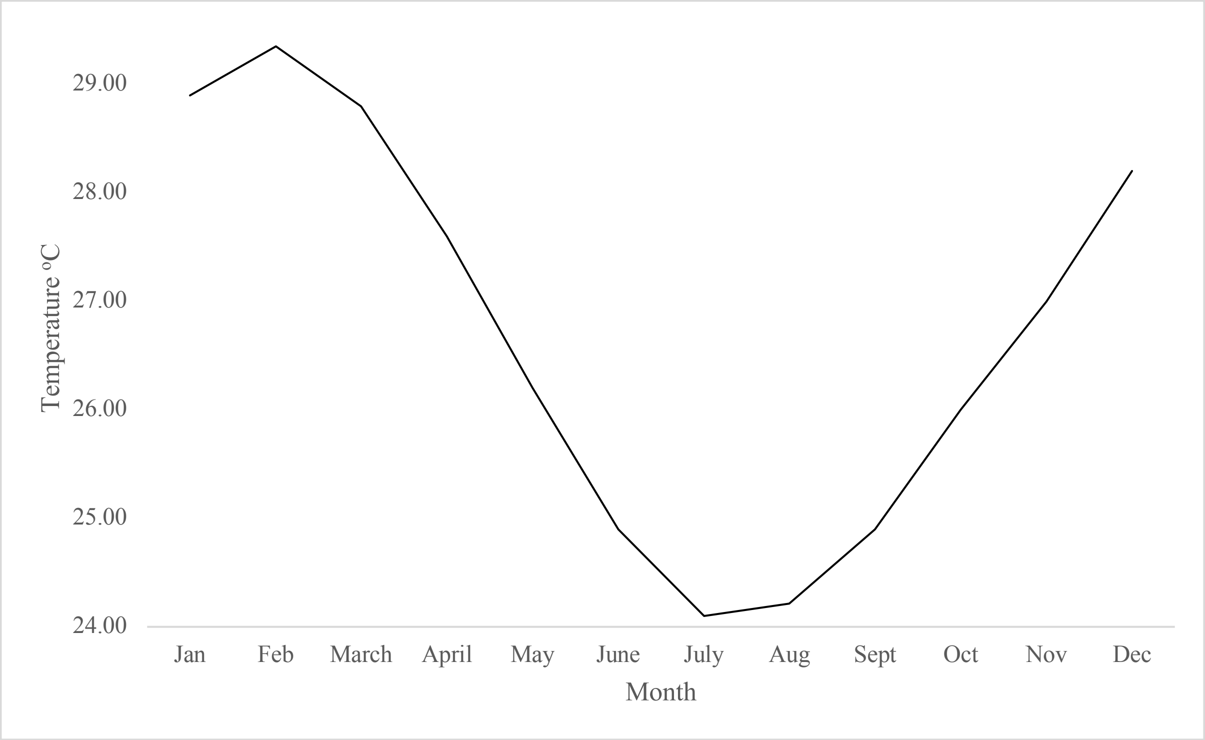


**Figure S7 Non-sequential eight-year sea water temperature average (1998 - 2017) from Moore Reef** (-16.8667°, 146.2334°')
(http://data.aims.gov.au/aimsrtds/datatool.xhtml?site=931&param=water%20temperature)

### Supplementary Tables

#### Table S1 Summary of materials and methods for the three experiments described in the manuscript.

| | **Ocean2100, Justus Liebig University Giessen** (Germany) | **The University of Hong Kong** (Hong Kong) | **Horniman Museum & Gardens, University of Derby** (United Kingdom) |
|---|---|---|---|
| **Experiment** | Menthol bleaching + Thermal performance experiment | Menthol bleaching + Symbiosis reestablishment with cultured Symbiodiniaceae | Completing gametogenic cycle and spawning *ex-situ* |
| **Colonies** | | | |
| Region of origin | Red Sea | Hong Kong | Australia (GBR) |
| Coordinates | 22.3054°, 38.9655° | 22.5308°, 114.3150° | -16.7000°, 146.0500° |
| Collection depth | 9 - 13 m | < 5 m | Unknown |
| No. of colonies | 3 | 4 | 35 |
| Type of biological replicates | Single polyps (< 1 cm) | Fragments (~ 3 cm) | Colonies (~ 10 cm) |
| **Rearing conditions** | | | |
| Tank size | 5 L | 6 L | 1000 L |
| System type | Closed | Closed | Closed |
| System description | Submersible pumps (Resun SP-500) + daily 10% water change | Submersible pumps (Atman AT301) + filters (Shiruba PF120 ) + 2x/week partial water change | 750 L broodstock tank with 250 L sump filtration. Ecoroll mechanical filter, protein skimmer and live rock for biological filtration. 10% system volume water change per month. |
| Temperature (°C) | 26 | 26 | ~24 - 29 (mimicking in situ conditions, see Fig. S8) |
| Light cycle (light:dark) | 12:12 | 12:12 | ~11:13 - ~13:11 mimicking in situ photoperiod (https://www.timeanddate.com/) |
| Light source | Fluorescent lamps (white) | Fluorescent lamps (white) | LEDs (White, Blue, Royal Blue, Green, Red, and UV) |
| Light intensity (µmol photons $m^{-2}$ $s^{-1}$) | 130 - 160 | 120 | mimicking in situ conditions (Craggs et al. 2017) |
| Salinity | 35 | 35 | 35 |
| Artificial sea salt | Aquarium Systems Reef Crystals | Instant Ocean Spectrum Brands | H2Ocean Pro reef salt |
| Corals maintained in filtered seawater | 1.2 µm | No | No |
| **Feeding** | | | |
| Type | Frozen adult Artemia | Powdered marine plankton (ReefRoids, Polyp Lab) | *Tisochrysis lutea*, *Chaetoceros calitrans*, *Rhodomonas salina*, newly hatched *Artemia* |

| | | | |
|---|---|---|---|
| Amount | 1 'shrimp'/polyp | 0.08g per 6L tank (calculated per manufacturer's recommendations) | *salina* napulii, frozen red plankton, rotifers and lobster / fish eggs |
| Frequency | Daily | Twice-weekly | Daily |

**Table S2 Relative abundance of cells distinguished as SymC, SymD or unlabelled from successfully FISH-hybridized samples**. Values are corrected for probe efficiency (92.3% ± 5.8 for SymC; 93.8% ± 4.0 for SymD)

| Sample | SymC (%) | SymD (%) | Unlabelled (%) |
|---|---|---|---|
| 1 | 77 | 4 | 19 |
| 4 | 42 | 19 | 39 |
| 5 | 82 | 17 | 1 |
| 7 | 79 | 9 | 12 |
| 8 | 78 | 17 | 5 |
| 11 | 47 | 50 | 4 |
| 12 | 41 | 54 | 5 |
| 14 | 37 | 63 | 1 |
| 16 | 59 | 20 | 21 |

**Table S3 Characterization of Symbiodiniaceae ITS2 sequences associated with colonies of *G. fascicularis* from Hong Kong**. Colonies collected from Crescent Island (Hong Kong) in March 2019. Sequencing depth: HK1_8 = 1,669, HK2_4 = 30.

| Colony | ITS2 sequence type (SymPortal output) | relative abundance (%) |
|---|---|---|
| HK1 | **C1** | 68.42 |
| HK1 | **C1c** | 15.40 |
| HK1 | C72k | 2.40 |
| HK1 | C3ju | 1.26 |
| HK1 | C1al | 1.20 |
| HK1 | C1ge | 1.02 |
| HK1 | C1b | 0.96 |
| HK1 | C1ch | 0.90 |
| HK1 | 281_C | 0.84 |
| HK1 | C1w | 0.60 |
| HK1 | C3 | 0.60 |
| HK1 | C3sa | 0.60 |
| HK1 | C1t | 0.54 |
| HK1 | C1bc | 0.54 |
| HK1 | 11561_C | 0.54 |
| HK1 | C3sf | 0.48 |
| HK1 | 73272_C | 0.42 |
| HK1 | C11 | 0.42 |
| HK1 | 25715_C | 0.42 |
| HK1 | 33564_C | 0.42 |
| HK1 | 6394_C | 0.42 |
| HK1 | 21512_C | 0.42 |
| HK1 | 21511_C | 0.42 |
| HK1 | 14089_C | 0.42 |
| HK1 | C42.2 | 0.36 |
| HK2 | **C1** | 80.00 |
| HK2 | **C1c** | 20.00 |

**Table S4 Summary of qualitative symbiosis reestablishment trial (one-time inoculum) on bleached *G. fascicularis* colonies from the Red Sea.** Donor colonies were used to produce cleaned Symbiodiniaceae extracts that were applied to the recipient colonies. Success rate denotes the number of polyps that successfully reestablished symbiosis from the total number of replicates used.

| Symbiont extract donor colony | Symbiont extract recipient colony | Success rate |
|---|---|---|
| RS1 | RS1 | 1/3 |
| RS2 | RS1 | 2/2 |
| RS3 | RS3 | 0/2 |

### Supplementary Text

#### Qualitative assessment of symbiosis reestablishment after menthol bleaching

We qualitatively explored the possibility of returning adult polyps to the symbiotic state after menthol bleaching using two approaches. Both of these employed bleached polyps previously used in the context of the thermal performance experiment, one month after the termination of the menthol bleaching treatment.

The first and simplest approach consisted of testing "environmental" symbiont acquisition. For this, 12 bleached polyps were transferred back to the long-term rearing aquarium system (Ocean2100 facility, JLU Giessen, Germany) and placed next to symbiotic colonies of *G. fascicularis* to be exposed to Symbiodiniaceae naturally released into the water column. Symbiont density in the aquarium seawater was not quantified, however their presence is expected based on published assessments of Symbiodiniaceae released by corals (e.g., Hoegh-Guldberg et al. (1987); Stimson and Kinzie (1991)).

The second approach consisted of inoculating seven bleached polyps with freshly isolated Symbiodiniaceae from symbiotic polyps. For this, a donor polyp from each colony was placed upside-down inside a 1.5 mL tube and centrifuged at 8,300 g for 1 min to detach the tissue from the skeleton (Wilkinson et al. 2016). The slurry was homogenized with 600 µl of FASW, centrifuged at 3,400 g for 1 min, and the resulting pellet resuspended twice to clean the algal fraction. Each donor (symbiotic) polyp was used to inoculate two to three polyps, from the same colony or from one respective other colony (Tab. S4). These received an equal amount of inoculum, however the concentration of algal cells was not quantified. For the inoculation, bleached polyps were placed in individual beakers, the Symbiodiniaceae inoculum was pipetted on the oral opening together with one Artemia shrimp, and allowed to feed for 3 h. Subsequently, inoculated polyps were transferred to the long-term rearing aquarium facility.

For both approaches, symbiosis reestablishment was visually assessed at irregular intervals in the following months, and photographically documented one year after (Fig. S5). Of the 12 colonies used for the environmental symbiont acquisition and of the 7 used for the tissue extract inoculation, 4 and 3 (30 and 43 %), respectively, successfully reestablished symbiosis and grew into small colonies that were indistinguishable from the symbiotic controls (Fig. S5).

<u>References</u>

Hoegh-Guldberg O, McCloskey LR, Muscatine L (1987) Expulsion of zooxanthellae by symbiotic cnidarians from the Red Sea. Coral Reefs 5:201–204

Stimson J, Kinzie RA (1991) The temporal pattern and rate of release of zooxanthellae from the reef coral Pocillopora damicornis (Linnaeus) under nitrogen-enrichment and control conditions. J Exp Mar Bio Ecol 153:63–74
